# Glimma: interactive graphics for gene expression analysis

**DOI:** 10.1101/096107

**Authors:** Shian Su, Charity W. Law, Casey Ah-Cann, Marie-Liesse Asselin-Labat, Marnie E. Blewitt, Matthew E. Ritchie

## Abstract

**Motivation:** Summary graphics for RNA-sequencing and microarray gene expression analyses may contain upwards of tens of thousands of points. Details about certain genes or samples of interest are easily obscured in such dense summary displays. Incorporating interactivity into summary plots would enable additional information to be displayed on demand and facilitate intuitive data exploration.

**Results:** The open-source *Glimma* package creates interactive graphics for exploring gene expression analysis with a few simple R commands. It extends popular plots found in the *limma* package, such as multi-dimensional scaling plots and mean-difference plots, to allow individual data points to be queried and additional annotation information to be displayed upon hovering or selecting particular points. It also offers links between plots so that more information can be revealed on demand. *Glimma* is widely applicable, supporting data analyses from a number of well established Bioconductor workflows *(limma, edgeR* and *DESeq2)* and uses D3/JavaScript to produce HTML pages with interactive displays that enable more effective data exploration by end-users. Results from *Glimma* can be easily shared between bioinformaticians and biologists, enhancing reporting capabilities while maintaining reproducibility.

**Availability and Implementation:** The *Glimma* R package is available from http://bioconductor.org/packages/devel/bioc/html/Glimma.html.

## Introduction

Analysis of gene expression via RNA-sequencing (RNA-seq) or microarray technologies produces large volumes of data that require highly condensed visual summaries to aid interpretation. Plots of summary statistics typically contain many thousands of points, and extracting details about particular genes or samples of interest can be non-trivial from these displays. Interactive graphics are highly effective tools for investigating data. They allow individual points to be queried, graphical filtering of data, the association of events between plots and animation of data, all of which can be useful for exploring large datasets. While there are mature packages for producing static plots in the R software environment, such as base *graphics* [1] and *ggplot2* [2], developments in interactive graphics are ongoing and to date very few Bioconductor packages enable effective interactive exploration of data. To solve this problem for the common task of gene expression analysis, we developed the *Glimma* package which adds interactivity to two popular data displays.

## Approach

*Glimma* is an R package that generates specialised interactive graphics to aid in the exploration of results from differential expression (DE) analyses. *Glimma*, which loosely stands for **interactive graphics from limma,** currently supports output from 3 popular Bioconductor [3] analysis workflows, namely from *limma* [4], *edgeR* [5] and *DESeq2* [6]. The main focus is on visualising results for **RNA**-seq and microarray DE analysis on gene-summarised counts. The plots generated by *Glimma* are those popularised by the *limma* package with additional interactive features inspired by the Degust [7] software.

The *Glimma* package uses the D3.js JavaScript library to create HTML pages with interactive plots and tables that have cross-interactions for exploring results of DE analysis. In contrast to Shiny [8] based interfaces that allow users to manipulate the data and plotting options, *Glimma* is intended purely for displaying the results of an analysis. It functions as a drop-in replacement for the calls to plotting functions commonly found in a *limma, edgeR* or *DESeq2* analysis pipeline. The resulting plots can be viewed and interacted with on any computer with a modern web browser.

The basic plots available in *Glimma* are the multi-dimensional scaling (MDS) plot available using the `glMDSPlot` function and mean-difference (MD) plot available with `glMDPlot`. The MDS plot is an unsupervised clustering technique based on the eigenvalue decomposition of euclidean distances between samples based on their gene expression profiles. The HTML output from `glMDSPlot` contains an MDS plot of two consecutive dimensions plotted against each other with each sample represented by a point in the display. The distance between two samples reflects the leading log-fold-change (logFC) or typical logFC of genes separating the samples. By default the top 500 genes are used to calculate distances unless specified otherwise. Alongside the MDS plot is a barplot that displays the proportion of variation in the data explained by each of the first few dimensions or eigenvectors in the MDS plot. Clicking on a bar in this panel will highlight two consecutive bars and display the associated dimensions in the MDS plot, while hovering over each of the points in the MDS plot brings up sample information such as sample labels and groups. Points may be colour-coded by experimental group information that the user may switch between in the case that there are multiple variables (e.g. genotype, sex, batch) to look for relationships. This display is useful for deciding on an appropriate model for the DE analysis, saving the need for making a series of static plots over several dimensions with different colour schemes in order to understand the important experimental variables to account for in the analysis.

The MD plot is used for identifying differentially expressed genes between two or more experimental conditions. The interactive MD plot output by the `glMDPlot` function contains three key components which interact with each other to show multiple aspects of the data in the one display. A screenshot of this output for a mouse RNA-seq dataset [9] provided with the package is shown in Figure 1. The main component is a plot of gene-wise summarised statistics which takes the top-left position of the HTML page. Gene-wise logFCs are plotted against gene-wise average log-expression values where each point on the plot represents a single gene. Differentially expressed genes can be highlighted in colour. Hovering or clicking on a gene (or point) within the main plot brings the expression of each sample for the selected gene into view in a second plot in the top-right position. In this plot, points can be stratified by either experimental factors or numeric values. At the same time, associated gene information is displayed in the table below. Users can scroll through the table looking for any gene of interest, or hone in on specific genes or groups of genes using the search bar above the table. A third function, `glXYPlot` enables customised summary plots, where other quantities besides the average log-expression and logFCs can be plotted. This can be useful for generating volcano plots or for comparing gene expression changes between two datasets. A series of RNA-seq and microarray use cases for *Glimma* are provided in the detailed Users’ Guide distributed with the package, and a further RNA-seq example is presented in Law *et al.* (2016) [10].

**Figure 1:**
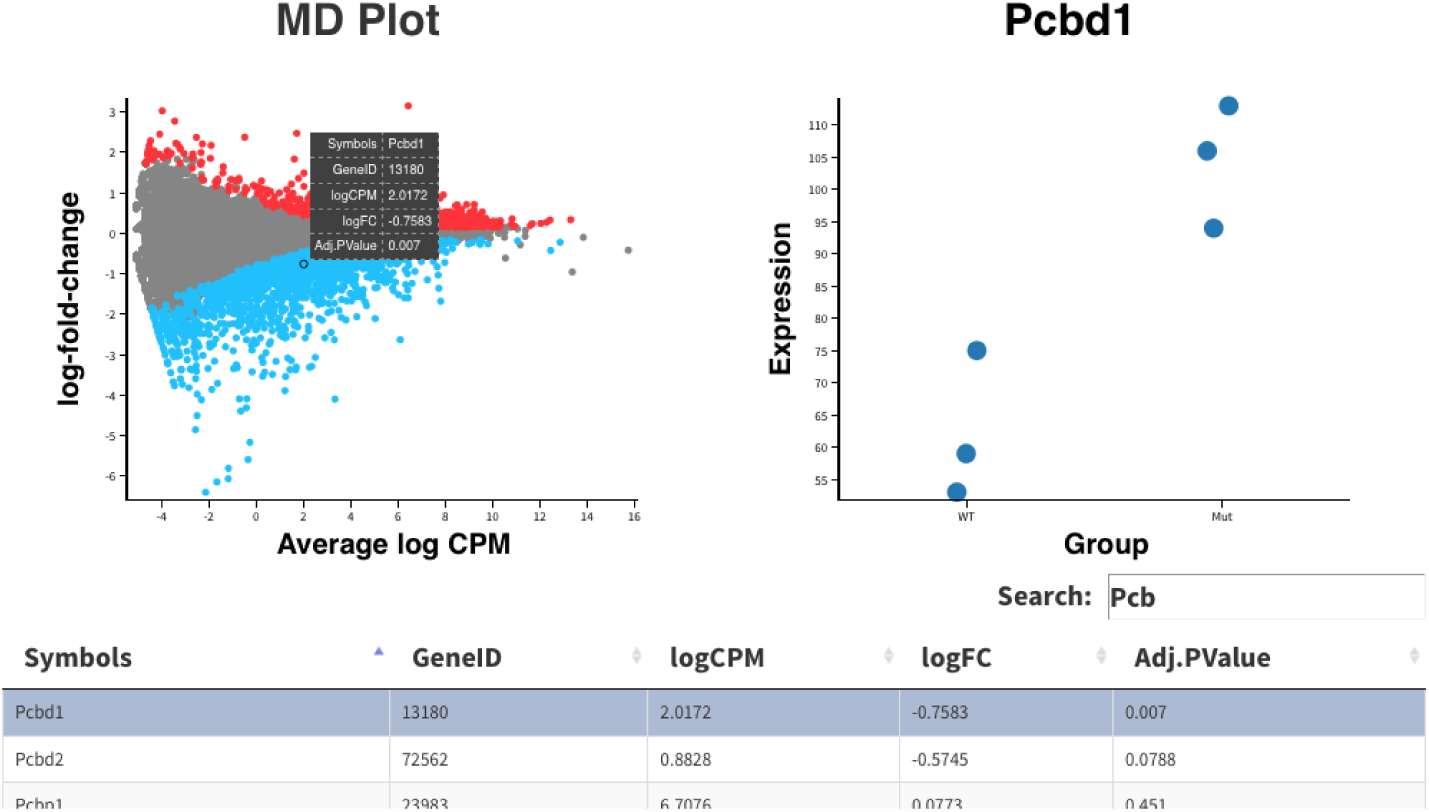
Screenshot of the HTML page generated by the `glMDPlot` function in *Glimma.* This visualisation combines the *limma*-style mean-difference plot (top-left) together with per sample expression information for a selected gene (*Pcbd1* in this case) stratified by sample type (top-right) and a table of differential expression results (bottom) with the same gene highlighted.

## Discussion

*Glimma* allows users to thoroughly and conveniently interrogate the results from a DE analysis by integrating several layers of gene-level and sample-level data in the one HTML output. Interactive *Glimma* graphics can be included in the analysis reports sent to collaborators to assist them in the prioritisation of genes for further study. Although the package was developed with gene expression data in mind, it can be used in any setting where there is a desire to connect summary-level data with sample-level observations and other annotations. Other uses include analysis of data from pooled genetic screens [11] and analyses of variability in single cell RNA-seq experiments and methylation array datasets. Future work includes adding new plotting capabilities, such as venn diagrams and heatmaps as optional displays and providing links to external databases to make it easier for users to obtain more annotation information. Replacement of the custom D3/JavaScript code that generates each interactive plot with functions from the rapidly developing *plotly* R package [12] is also planned to improve maintainability.

## Acknowledgements

We thank Dr Ahmed Mohamed for advice on implementation and Dr Mike Love for helpful feedback and example *DESeq2* code.

